# Topological analysis of brain dynamics in autism based on graph and persistent homology

**DOI:** 10.1101/2022.05.14.491959

**Authors:** Alireza Talesh Jafadideh, Babak Mohammadzadeh Asl

## Abstract

Autism spectrum disorder (ASD) is a heterogeneous disorder with a rapidly growing prevalence. In recent years, the dynamic functional connectivity (DFC) technique has been used to reveal the transient connectivity behavior of ASDs’ brains by clustering connectivity matrices in different states. However, the states of DFC have not been yet studied from a topological point of view. In this paper, this study was performed using global metrics of the graph and persistent homology (PH) and resting-state functional magnetic resonance imaging (fMRI) data. The PH has been recently developed in topological data analysis and deals with persistent structures of data. The structural connectivity (SC) and static FC (SFC) were also studied to better show the advantages of DFC analysis. Significant discriminative features between ASDs and typical controls (TC) were only found in states of DFC. Moreover, the best classification performance was offered by persistent homology-based metrics in two out of four states. In these two states, some networks of ASDs compared to TCs were more segregated and isolated (showing the disruption of network integration in ASDs). The results of this study demonstrated that topological analysis of DFC states could offer discriminative features which were not discriminative in SFC and SC. Also, PH metrics compared to graph global metrics can open a brighter avenue for studying ASD and finding candidate biomarkers.

**Highlights:**

1. States of dynamic functional connectivity (DFC) were more informative than static FC and structural connectivity when comparing ASDs with controls.
2. Global metrics of persistent homology (PH) in comparison to graph ones could better distinguish between ASDs and controls.
3. The PH metrics could offer the best classification performance in dynamic states where the networks of ASDs compared to controls were more segregated and isolated.

## 1. Introduction

Autism spectrum disorder (ASD) is a heterogeneous neurodevelopmental disorder mainly characterized by deficits in social communication and interaction, restricted interests, and repetitive behaviors or activities (American Psychiatric Association, 2013). Unfortunately, ASD is a heterogeneous disorder with a high-speed prevalence (Elsabbagh et al., 2012). Hence, reliable and early diagnosis has been a very vital demand and many kinds of research have been carried out to answer it by finding biomarkers and features (Goldani et al., 2014; Woo and Wager, 2015; Drysdale et al., 2016; Li et al., 2017). It has been shown that the brain structure and function are affected by this disorder (Ha et al., 2015). As a result, neuroimaging technologies and techniques can be employed to find quantifiable features, and consequently reliable biomarkers for ASD diagnosis using the brain structural and functional changes (Grecucci et al., 2017).

Two of the most popular techniques for analyzing the brain behavior and changes are the structural connectivity (SC) and functional connectivity (FC), in which the relations between brain areas (regions of interest, ROIs) are estimated (Uddin et al., 2013; Ha et al., 2015; Mash et al., 2018). The resting-state functional magnetic resonance imaging (rfMRI) data is popular for FC computation, due to its fast and task-free nature (Biswal, 2012; Hull et al., 2017). The conventional static FC (SFC) and newly developed dynamic FC (DFC) are two common methods for the computation of FC. The SFC computes FC using all time points of functional signals. The SFC studies offered both stronger (over) and weaker (under) connectivity for the ASD group compared to TC one (Cheng et al., 2015; Iidaka, 2015; Hull et al., 2016). This contradictory behavior of over- and under-connectivity may be originated from overlooking the transient behavior of fMRI data (Mash et al., 2019). To overcome this problem, the DFC employs the sliding window (SW) technique and computes a sequence of FC matrices using a subset of total time points. Then, these matrices are clustered into different FC “states” of the brain (Preti et al., 2017; Rashid et al., 2018; Aggarwal and Gupta, 2019; Mash et al., 2019). It has been stated that DFC might provide answers to SFC conflict results by residing weaker and stronger connectivities in different states (Mash et al., 2019). Mash and colleagues (Mash et al., 2019) reported the reduction of the default mode network segregation. This reduction happened by involving the somatomotor network in one state and fronto-parietal and executive networks in another state. Rashid and colleagues (Rashid et al., 2018) reported the positive connection between the dwelling time and the severity of autistic traits’ in a “globally disconnected” state. Less switching frequency between states of ASDs has also been reported, suggesting insufficient dynamic behavior of the brain (de Lacy et al., 2017; Fu et al., 2018; Watanabe & Rees, 2017).

The SC is another type of connectivity, in which a bundle of fibers connects the brain ROIs. The diffusion tensor imaging (DTI) tractography is usually used to rebuild the fiber tracts, which in turn are employed to compute the strength of the connections, and consequently estimate the SC matrix (Mukherjee and McKinstry, 2006; Jou et al., 2011; Mash et al., 2018). Using the DTI data, increased mean diffusivity (MD) in the whole frontal lobe (Sundaram et al., 2008), a temporal portion of the left superior longitudinal fasciculus (Nagae et al., 2012), and thalamo-cortical connections (Nair et al., 2013) were found when studying children with ASD.

The human brain is inherently complex, which in turn results in complex architecture for brain connectivity (Bullmore and Sporns, 2009). To handle this complexity, some approaches have been developed to study connectivities from a topological point of view. One of these approaches is graph theory, simplifying the connectivity architecture by a graph whose vertices are ROIs and edges represent the FC between ROIs (De Vico Fallani et al., 2014; Song et al., 2019). The most commonly used metrics of the graph are categorized into local and global ones (Fornito et al., 2016; Farahani et al., 2019). The earlier ones describe the network behavior at the ROI level and the latter ones describe the properties governing the entire brain network.

The ASD leads to abnormal FCs (Mash et al., 2019; Rashid et al., 2018), and consequently abnormal values of graph metrics. Therefore, these abnormalities can be revealed and considered as candidate biomarkers (Hull et al., 2016). Di Martino and colleagues (Di Martino et al., 2013) reported that ASDs compared to typical controls (TCs) had more connections (degrees) in the cortical and subcortical ROIs. Redcay E and colleagues (Redcay E et al., 2013) investigated the right parietal region of the default mode network and found a higher betweenness centrality (a metric indicating the ROI centrality) for ASDs. Both of these findings represent the over-connectivity in ASDs (Hull et al., 2016). Rudie and colleagues (Rudie et al., 2013) found that adolescents with ASD had more global and less local efficiency than TCs. By studying adult ASDs in comparison to adult TCs, Itahashi and colleagues (Itahashi et al., 2014) reported characteristic path length and clustering coefficient reduction. These results state that the randomness of ASDs’ networks is more than TCs (Rudie et al., 2013; Keown et al., 2017). The analysis of Keown and colleagues (Keown et al., 2017) offered the globally decreased cohesion and increased dispersion of networks. Cohesion quantified how well nodes from a normative network grouped within the community structure. Dispersion quantified how distributed nodes from a network were in the community structure. These results showed that the segregation and integration of ASD networks were impaired (Keown et al., 2017). Altogether, impaired functional organizations of the autistic brain were found in both local and global metrics of the graph (Farahani et al., 2019).

Another approach for characterizing the topological organization of brain connectivities is using the persistent homology (PH) concept and metrics (Edelsbrunner snd Harer, 2010; Zomorodian et al., 2005). This technique has been recently developed to detect structures of brain networks persistent across multiple scales (multiple graph filtration values) (Lee et al., 2012; Chung et al., 2015; Yoo et al., 2017; Lee et al., 2017; Xing et al., 2022). These structures are components, holes, voids, and higher dimensions polytopes. The topological structures with a shorter duration (lower persisting) are considered noise, and the structure with a longer duration (higher persisting) represents an essential structure. Based on the birth and death values, the number of studied structure (for example holes), and filtration values, many different quantitative metrics of PH can be defined to characterize the network. PH method in comparison to the graph theory method can better reveal topological changes and avoid the problem of scale threshold selection (applying threshold on connectivity matrix for keeping the strong connections) (Otter et al., 2017; Xing et al., 2022).

The PH technique has been successfully applied to study the brain network structure of Alzheimer disease (AD) (Kuang et al., 2019a,b, 2020a,b), epilepsy (Choi et al., 2014), autism spectrum disorder, attention-deficit hyperactivity disorder (Lee et al., 2012, 2017), etc. Kuang and colleagues (Kuang et al., 2019a) proposed the integrated persistent feature (IPF) plot to analyze the rfMRI data of AD, mild cognitive impairment (MCI), and normal control (NC) groups. To compute IPF values, both the number of components and filtration values were employed. The IPF plot was attained by plotting the IPF values with respect to filtration values. The slope of the IPF curve could describe the spatial dynamics of the brain network. Using the slope values, the authors reported significant differences between AD and MCI and between AD and normal control (NC) groups. The IPF curves are usually nonlinear. Therefore, the slope of IPF may not appropriately characterize the behavior of persistent features. To consider this concern, Kuang and colleagues (Kuang et al., 2019b) used the kernel technique to extract a kernel-based feature from the IPF curve. By studying the fluorodeoxyglucose positron emission tomography (FDG-PET) imaging data, they found that their proposed feature, in contrast to the slope feature, could offer a significant difference between all pairs of (AD, MCI, NC). It was also found that the IPF-based features were more robust and discriminator than graph metrics (Kuang et al., 2019a,b, 2020a,b). Lee and colleagues (Lee et al., 2017) computed the Betti numbers for components using the single linkage matrix (SLM) (Lee et al., 2012) (the number of components, holes, voids, etc are also called the Betti number). The authors analyzed the asymmetric change in the brain network using the PET and MRI data. They found that the brain networks of ASD, attention deficit hyperactivity disorder (ADHD) children, and controls differ, with ASD and ADHD showing asymmetrical changes of connected structures between metabolic and morphological connectivities.

In this study, the SC, SFC, and states of DFC of ASD and TC groups are studied from a topological point of view using different global metrics of graph and PH. To be revealed the discriminative capacity of each connectivity type, they are analyzed separately. To the best knowledge, this is the first attempt at topological analysis of ASD dynamic states using graph and PH metrics. Also, SC and SFC of ASDs have not been yet studied using PH metrics. We aim to know 1) which connectivity type can be more informative for discriminating between ASDs and TCs?; 2) which states of DFC can offer more discriminative features?; 3) can PH global metrics be more discriminative than graph ones?. For evaluation purposes, statistical analysis and classification are used.

The rest of the paper is organized as follows: Section 2 describes the dataset, preprocessing methods, and brain ROIs and networks. Then, the computation methods of SC, SFC, and states of DFC are given. This section is continued by defining the graph and PH metrics. The last two subsections of this section are devoted to statistical analysis and classification evaluations. Section 3 provides the results. The discussion about the results and conclusions are respectively given in sections 4 and 5.

## 2. Materials and methods

A schematic of the experimental design is presented in Figure 1.

**Figure 1.**
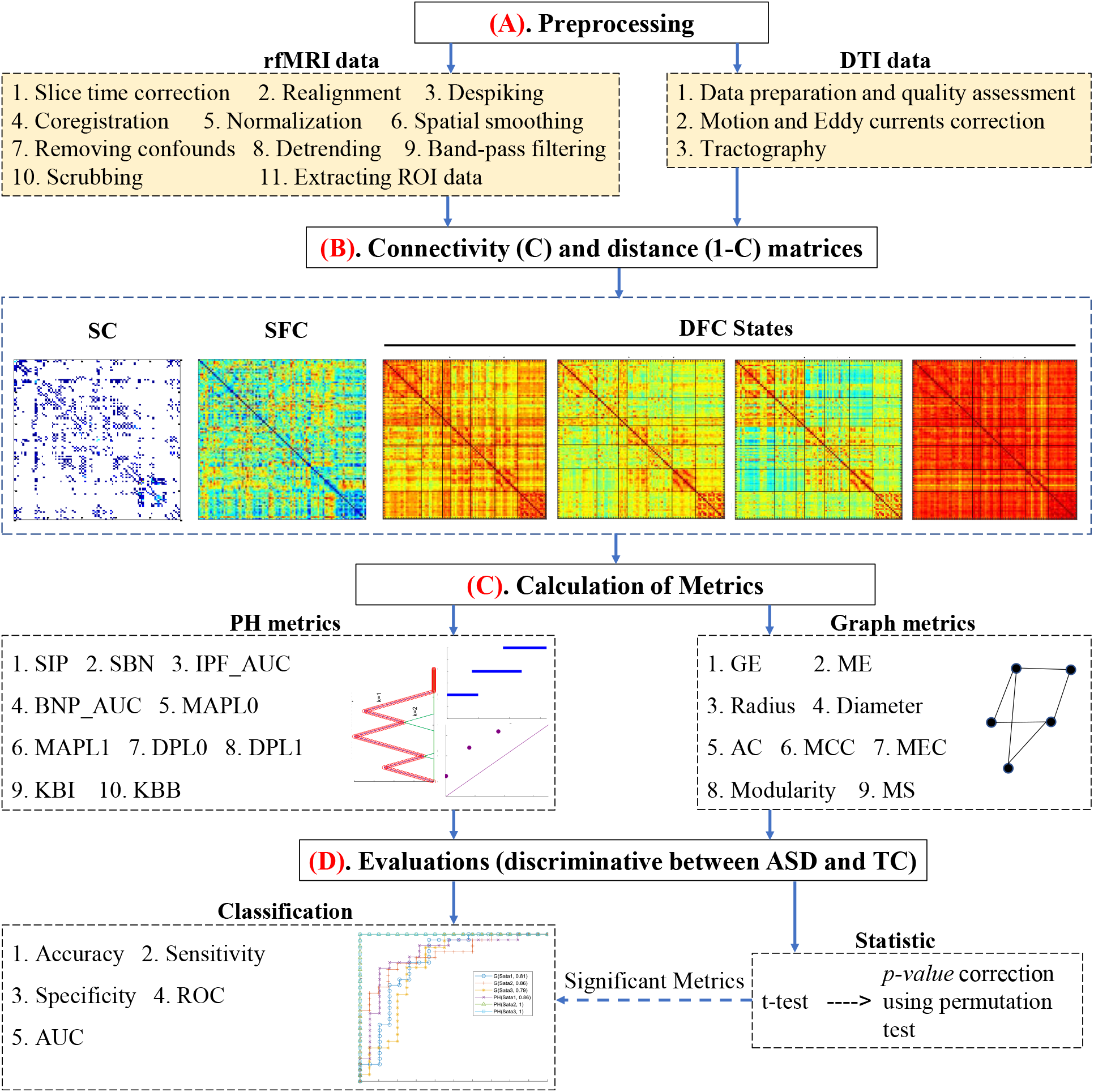
Experimental design. SC, SFC, and DFC stand for structural, functional, and dynamic functional connectivity, respectively. The panels **(A-C)** are performed for ASD and TC groups, separately. The panels **(C)** and **(D)** are performed for each type of connectivity, separately. In panel **(D)**, the ability of different metrics and connectivity types in discriminating between ASD and TC subjects are examined. For classification, only significant metrics and connectivity types with significant metrics are used.

### 2.1. Participants and data acquisition

The SDSU of ABIDE II database was employed in this study. Both ASD and TC groups of this dataset had mostly male subjects. Accordingly, only the male subjects were analyzed. Due to the problem of motion artifact, one TC subject with ID number “28852” was ignored, (see section 2.2.1). Two groups were matched on age, handedness, and head motion while showing a significant difference in the social responsiveness scale (SRS). The SRS value of one ASD subject was missing. The sample characteristics are listed in Table 1. The framewise displacement (FD) is explained in subsection 2.2.1.

**Table 1.** Sample characteristics.

|  | ASD | Control | Statistic | *p* |
| --- | --- | --- | --- | --- |
| Age (year) | 12.61±3.3<br>[7.4-17.8] | 13.55±3.1<br>[8.1-17.7] | $t(46)=-1.01$ | 0.32 |
| Sex | 26 Male | 22 Male | - | - |
| Handedness | 21 Right | 18 Right | $\chi^2(1)=0.01$ | 0.92 |
| SRS total | 82.56±9.41<br>[62-94] | 42.6±5.37<br>[35-52] | $t(45)=17.54$ | **<1e-15** |
| rFMRI FD (mm) | 0.04±0.03<br>[0.01-0.12] | 0.04±0.03<br>[0.01-0.11] | $t(46)=0.85$ | 0.4 |
| # of rFMRI time points with<br>FD>0.5mm | 0.9±2.5<br>[0-11] | 0.23±0.87<br>[0-4] | $t(46)=0.66$ | 0.51 |
| DTI FD (mm) | 0.27±0.09<br>[0.11-0.52] | 0.25±0.08<br>[0.18-0.55] | $t(46)=0.21$ | 0.83 |
| # of DTI time points with<br>FD>0.5mm | 4.9±5.2<br>[0-16] | 3.91±5.38<br>[0-22] | $t(46)=1.24$ | 0.22 |

Abbreviations: AAL = Automated Anatomical Labelling; DTI = Diffusion tensor imaging; rFMRI = rest functional magnetic resonance imaging; FD = framewise displacement; SRS = social responsiveness scale.

In this paragraph, the data collection information of SDSU is represented. The data was collected using GE 3T MR750 scanner with eight-channel head coils. High-resolution structural images were acquired according to SPGR standards’ T1-weighted sequences and with TR/TE of 8.136/3.172 ms, flip angle of 8°, the field of view (FOV) of 256×256 mm^2^, resolutions of 1 mm^3^, and the number of slices of 172. The rfMRI dataset was collected using the standard gradient echo-planar imaging (EPI) sequences with TR/TE of 2000/30 ms, flip angle of 90°, FOV of 220×220 mm^2^, matrix size of 64×64, pixel spacing size of 3.4375×3.4375 mm^2^, slice thickness of 3.4 mm, slice gap of 0 mm, axial slice of 42, and total volumes of 180. The DTI protocol was based on the EPI sequence. The DTI images consisted of 61 weighted diffusion scans with a *b* value of 1000 sec/mm^2^ and one unweighted diffusion scan with a *b* value of 0 sec/mm^2^. The other parameters of data recording included TR/TE of 8500/78 ms, FOV of 128×128 mm^2^, slice in-place resolutions of 1.875×1.875 mm^2^, slice thickness of 2 mm, and the number of axial slices of 68.

### 2.2. Data preprocessing

#### 2.2.1. rfMRI preprocessing and time series extraction

The SPM (http://www.fil.ion.ucl.ac.uk/spm/) and AFNI (https://afni.nimh.nih.gov) software packages and personal codes were used to preprocess the rfMRI data. To avoid the adverse effect of T1-equilibration, the first five volumes were removed. The typical preprocessing steps were carried out for each subject. These steps were slice-timing correction using the middle slice as the reference time frame, motion correction, despiking using the AFNI’s 3dDespike, coregistration of T1 image and functional images using the spm_coreg function, warping images to the standard Montreal Neurological Institute (MNI) space, and spatial smoothing using a Gaussian kernel with a 5-mm full width at half maximum (FWHM = 5 mm). The MNI space in SPM12 was defined by a template created by nonlinear registration of 152 T1-weighted images.

For more noise reduction, some post-preprocessing steps were performed. Fourteen confounds were regressed out from all voxels’ time series. These confounds were six motion parameters and their first temporal derivatives, one principal component of cerebrospinal fluid (CSF), and one principal component of white matter (WM) signals. Using the mask of the CSF/WM, available in WFU PickAtlas software (http://fmri.wfubmc.edu/software/pickatlas), the time series of its voxels were extracted. Then, principal component analysis (PCA) was employed to compute the principal components of CSF/CM. For each of the CSF and WM, one dominant component took more than 99% variance of the time series. Thus, they were selected as confounds (Caballero-Gaudes and Reynolds, 2017). In the next steps of preprocessing, the time series signals were cleaned more by removing the linear, quadratic, and cubic trends, and then, applying a band-pass fifth-order Butterworth filter (0.01-0.1 Hz).

Some of the volumes might be corrupted due to motion artifacts. The differences between motion parameters of two consecutive time points (a total of six differences for six motion parameters) were calculated to detect the corrupted volumes. Then, the FD was computed using the square root of the sum of squares of computed differences. To transform the radians of rotational parameters to the millimeter, the head was regarded as a sphere of radius 50 mm (Power et al., 2012). The time points with FD >0.5 mm and the two subsequent time points were censored (Mash et al., 2019). As with Mash and colleagues (Mash et al., 2019), time series fragments with <10 consecutive time points remaining after censoring were also excluded. After censoring corrupted time points, if the number of remaining time points of a subject was less than 75% of the number of the initial time points (175), that subject was rejected to be used in this study. The TC subject with ID = 28852 was rejected. The average of eliminated time points for ASD and TC were 0.9 and 0.41, respectively. Totally, 23 subjects of ASD and 19 subjects of TC had no censoring time points, i.e., no significant difference between the two groups (χ2 (1) = 0.048, *p* = 0.82). The maximum censored time points for one subject of ASD and TC were 19 and 16, constituting 34.54% and 84.21% of the total number of censored points of these groups, respectively. Two groups did not show a significant difference in terms of the number of eliminated time points (t(55) = 0.82, *p* = 0.41). It is more convenient to have equal numbers of time points for all understudy subjects. Hence, given that the maximum number of censored time points and the initial time points were respectively 19 and 175, the first 156 time points of each subject were selected for further analysis (Goldani et al., 2014).

#### 2.2.2. DTI preprocessing

In this study, the ExploreDTI software was used to preprocess the DTI data and compute the structural connectivity matrix. Initially, data preparation and quality assessment were performed in ExploreDTI. Then, subjects’ motion and Eddy currents of diffusion data were corrected. After that, tractography was carried out using deterministic fiber tracking. The FDs of DTI data were computed, which are summarized in Table 1. In ExploreDTI, outliers are detected and removed from the tensor estimation procedure using the REKINDLE (Robust Extraction of Kurtosis INDices with Linear Estimation) (Tax C.M.W. et al, 2015).

### 2.3. Brain ROIs and networks

In this study, the brain was parcellated by Schaefer atlas (Schaefer et al., 2018) with 100 ROIs. This atlas, which is a functional atlas, respects the boundaries of the 7 Yeo-Krienen functional atlas (Yeo et al., 2011). In this atlas, the cerebral ROIs are assigned to 7 and 17 networks. The atlas with seven networks was used in the current study. These networks included default mode network (DMN), control network (CN), limbic network (LN), salient/ventral attention network (SVAN), dorsal attention network (DAN), somatomotor network (SMN), and visual network (VN). Hereinafter, this functional atlas with 100 ROIs is denoted by FA100.

For each ROI, one signal was obtained by averaging the time series of that ROI voxels. The final rfMRI data was 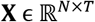 where *N* = 100 and *T* = *156* were the numbers of ROIs and time samples, respectively.

### 2.4. SC, SFC, DFC, and distance matrices

By using the tractography results and atlas information, the structural connectivity matrix was computed for each subject. In this matrix, the strength of connectivity between the two regions was equal to the number of tracts existing between them. In the next step, each row of the matrix was divided by the sum of its elements. By doing so, the value of entry (*i,j*) expressed the connectivity probability from region *i* to region *j*, which was not equal to that from region *j* to region *i*. Thus, the connectivity matrix, also called the adjacency matrix **A**, was not symmetric. As the final step, the symmetric adjacency matrix **A’** was obtained as **A’** = (**A** + **A^T^**)/2 where.^T^ denotes the transpose operator.

The SFC was computed between ROIs using Pearson correlation to have one static connectivity matrix for each subject.

To consider the transient behavior of FC, the DFC was employed through the SW technique. In SW, a sequence of windows with one TR shift was applied to the time series of each ROI. The window was created by convolving a rectangle (width of 22 TRs) with a Gaussian (standard deviation of 3 TRs) (Allen et al., 2014). Then, one FC matrix was computed for each window. These transient FC matrices of all subjects were clustered using the k-means with 100 times repetition and a maximum iteration of 1000. (Vassilvitskii, 2006). The elbow metric, which is the ratio of the within-cluster sum of squared distances (WCSSD) to between clusters SSD (BCSSD), was employed to find the optimum number of clusters. In WCSSD, the SSD of all members of a given cluster was calculated from the center of the cluster, and this process was repeated for all clusters. Finally, the sum of all SSDs was considered the WCSSD. In BCSSD, for a given cluster, the SSD of all transient FCs, which were not members of the cluster, was computed from its center, and this process was repeated for all clusters. Finally, the sum of SSDs was considered the BCSSD. After clustering, the clusters’ centers and members respectively represented the DFC’ states and states’ members.

ASD and TC groups may have a different number of states. To consider this, the clustering process was carried out for ASD and TC groups, separately. The closest TC state to a given ASD state was specified using the Euclidean distance so that the final matched states were the best matching pairs. In each state, the median of all FC matrices belonging to a given subject was taken to have one FC matrix for that subject.

For each subject, the output of SC, SFC, and states of DFC methods was a connectivity matrix **C**. However, one distance matrix was needed to calculate the PH metrics and some metrics of the graph. In this study, this matrix was attained through **1-C**, where **1** is an all-one matrix.

It should be noted that the SC, SFC, and each state of DFC were separately investigated for finding significant discriminative features.

### 2.5. Graph metrics

A graph is a set of vertices and edges. The connectivity matrix can be modeled by a graph in which the ROIs are vertices, and the connectivity strengths are the weights of edges. This modeling helped us to investigate topological differences between ASD and TC groups using graph metrics. In the following, some of the graph global metrics are defined (Fornito et al., 2016).

#### Global efficiency (GE)

GE is equal to the average inverse shortest path length in the network. In this study, the shortest path between two ROIs is defined as the is the distance between them.

#### Mean Eccentricity (ME)

For each ROI, the Eccentricity is equal to the maximum distance between that ROI and the rest of ROIs. ME equals the average eccentricity of all ROIs.

#### Radius

The minimum value of eccentricity is equal to the radius.

#### Diameter

The maximum value of eccentricity is equal to the diameter.

#### Assortativity coefficient (AC)

Each connection involves two ROIs; one starts and the other ends the connection. Let us call the degrees of the first and second involved ROIs x and y, respectively. By doing this, for all available connections, we obtain two vectors X and Y, the first of which has a set of degrees x and the second a set of degrees y. By calculating the correlation coefficient between X and Y, the value of AC is obtained. This coefficient takes values between −1 and 1. The positive values show that ROIs with similar degrees tend to connect. The negative AC value states that the tendency of ROIs with larger degrees is to connect to ROIs with smaller degrees.

#### Mean of clustering coefficient (MCC)

The Clustering coefficient is the fraction of triangles around an ROI and takes values between 0 and 1. The value 1 means that connected ROIs to a given ROI are also connected. Fewer connections in the neighborhood of a given ROI lead to less value of the clustering coefficient. In this study, the mean of clustering coefficient values over ROIs was considered for each subject.

#### Mean of eigenvector centrality (MEC)

Connections originating from high-scoring ROIs contribute more to the score of the ROI in question than connections from low-scoring ROIs. The EC of an ROI expresses the impact of the ROI on a network. ROI with high EC connects to ROIs with high scores. In this study, the mean of EC values over ROIs was considered for each subject.

#### Modularity

This metric expresses a network how well has been divided into groups of ROIs. In a network with high modularity, there exist dense connections within the groups and sparse connections between the groups of ROIs.

#### Mean of strength (MS)

The strength of a given ROI is the sum of weights of adjacent edges to that ROI. In this study, the mean of strength values over ROIs was considered for each subject.

All these metrics were computed using the codes of the brain connectivity toolbox (Rubinov and Sporns, 2010).

### 2.6. PH

Here we are interested in homology, which associates one vector space H_i_(X) to a space X for each natural number *i* ∈ {0,1,2,… }. The H_i_ is also called the i^th^ homology group. The dimension of H_0_(X) counts the number of components in X, the dimension of H_1_(X) is a count of the number of holes, and the dimension of H_2_(X) is a count of the number of voids (Otter et al., 2017). The Betti number *β_i_* represents the dimension of H_i_ (panel (A) of Figure 2).

**Figure 2.**
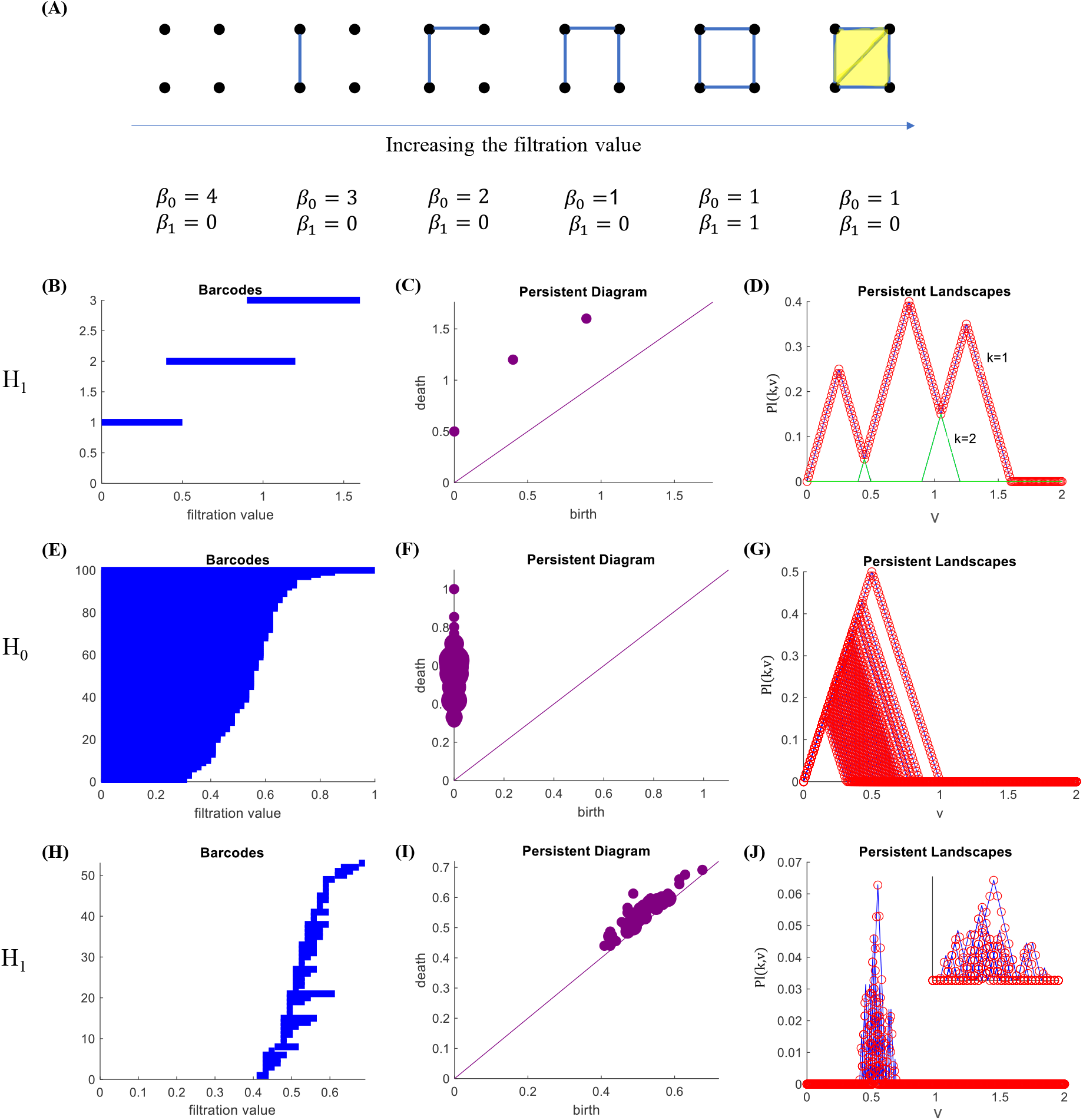
Examples of Betti numbers, barcodes, persistent diagrams, and persistent landscapes. **(A)** The connection between points and consequently the Betti numbers *β*_0_ and *β*_1_ change when increasing the filtration value. **(B, C, D)** For simple illustration purpose, the barcodes, persistent diagrams, and persistent landscapes for holes with [birth, death] of [0,0.5], [0.4,1.2], and [0.9,1.6] are plotted. **(E, F, G)/(H, I, J)** The plots of the second row are repeated for components/holes of SFC of one TC subject. In panel (J), a zoomed-in view plot is also shown. The blue lines of (**D, G, J**) represent the triangles corresponding to each barcode. These triangles are the fundamental elements of persistent landscapes.

The PH is a method for computing topological features that persist across multiple scales. As a result, such features are expected to provide real data properties rather than noise, perturbation, or inappropriate parameter selection (Carlsson, 2009). To find the PH of a space, one must first represent the space as a simplicial complex. A simplicial complex is made up of a set of points, edges, triangles, tetrahedra, and higher dimensions polytopes. A simple example of the simplicial complex is shown in panel (A) of Figure 2.

Suppose that each of the ROIs is located at a point in Euclidean space. By applying the filter to the distance values existing between ROIs, several ROIs with distances less than the filter value are connected. By increasing the filtration value, the number of connections increases, and the resulting simplicial complex would have more components, edges, triangles, etc (panel (A) of Figure 2). Each component or hole or void is generated for a certain amount of filtration value, and lasts by increasing the amount of filter, and after reaching a certain amount of filter will die due to the connections that have occurred. Barcodes can be used to show the persistence of a component, hole, or void (panels (B, E, H) of Figure 2). In addition, the persistent diagram can be used, the horizontal and vertical axes of which represent the birth and death of a component, hole, or void, respectively (panels (C, F, I)).

In this study, the components and holes of the subjects’ distance matrices were examined. The Javaplex-4.3.4 software was used to extract the components and holes and their birth and death (Adams et al., 2014).

#### 2.6.1. Persistent landscape

One of the main challenges of working with PH is combining barcodes or PDs with statistics and machine learning. This is because the space of barcodes lacks geometric properties (Otter et al., 2017). An alternative approach is using the persistence landscape developed by Peter Bubenik (Bubenik, 2015). The basic idea is to convert barcodes to a function of a variable such as v. By considering that the statistics related to working with functions have been developed for a long time, then statistical analysis can be performed using these functions. The persistent landscape (PL) is defined for each of the homology groups, i.e., component and holes, separately. Suppose we have up to n barcodes with the birth and death values of [b_j_, d_j_] where j = 1,2,…, n. The PLs are a set of pl (k, v) functions which are defined as follows:

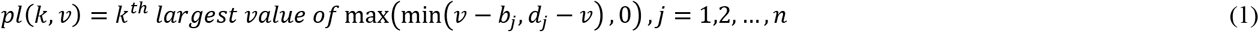

In this study, according to the minimum and maximum values of birth and death values, the value of v was defined from 0 to 2 with a step of 0.01. Thus, the maximum value of k can be n, meaning there are at most n PL functions. Examples of PLs are shown in panels (D, G, J) of Figure 2.

#### 2.6.2. IPF and Betti number plot

The minimum spanning tree (MST) of the distance matrix is needed to define the IPF and Betti number plot (BNP) (Kuang et al., 2019a). The MST is a subset of the edges of a connected graph that connects all the ROIs, without any cycles and with the minimum possible total edge weight (Lee et al., 2012). In this study, the Single linkage dendrogram was used for computing the MST (Lee et al., 2012). For M ROIs, there are M-1 weights of MST which are sorted from small to large to form the filtration values *λ* = {*λ*_0_ = 0 < *λ*_1_,… < *λ*_*M*-1_}. Using these values, the BNP and IPF are defined as

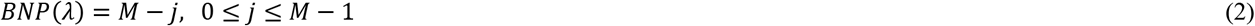

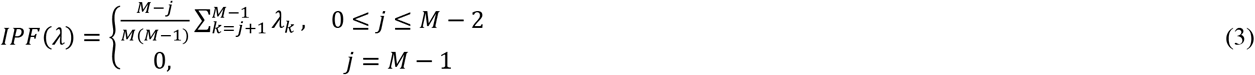

It should be noted that BNP and IPF are constructed using the MST which has no cycles (no holes). Therefore, these two criteriums can only be used to analyze components of a tree. As it is clear from the BNP and IPF equations, the former only counts the number of components while the latter deals with the filtration values. The IPF and BNP gradually decrease, and one fully connected component is constructed eventually when filtration values increase from 0 to 1.

The results of BNP and IPF for the TC and ASD groups are shown in Figure 3. For each group, these results were obtained by averaging the BNP and IPF of all subjects. The IPF plots show more differences between ASD and TC groups compared to the BNP plot. The least/the most differences between ASD and TC subjects are seen in SC/States2-3 of DFC. On average, the slop of IPF changes in State2/State3 is more/less negative for ASDs compared to TCs.

**Figure 3.**
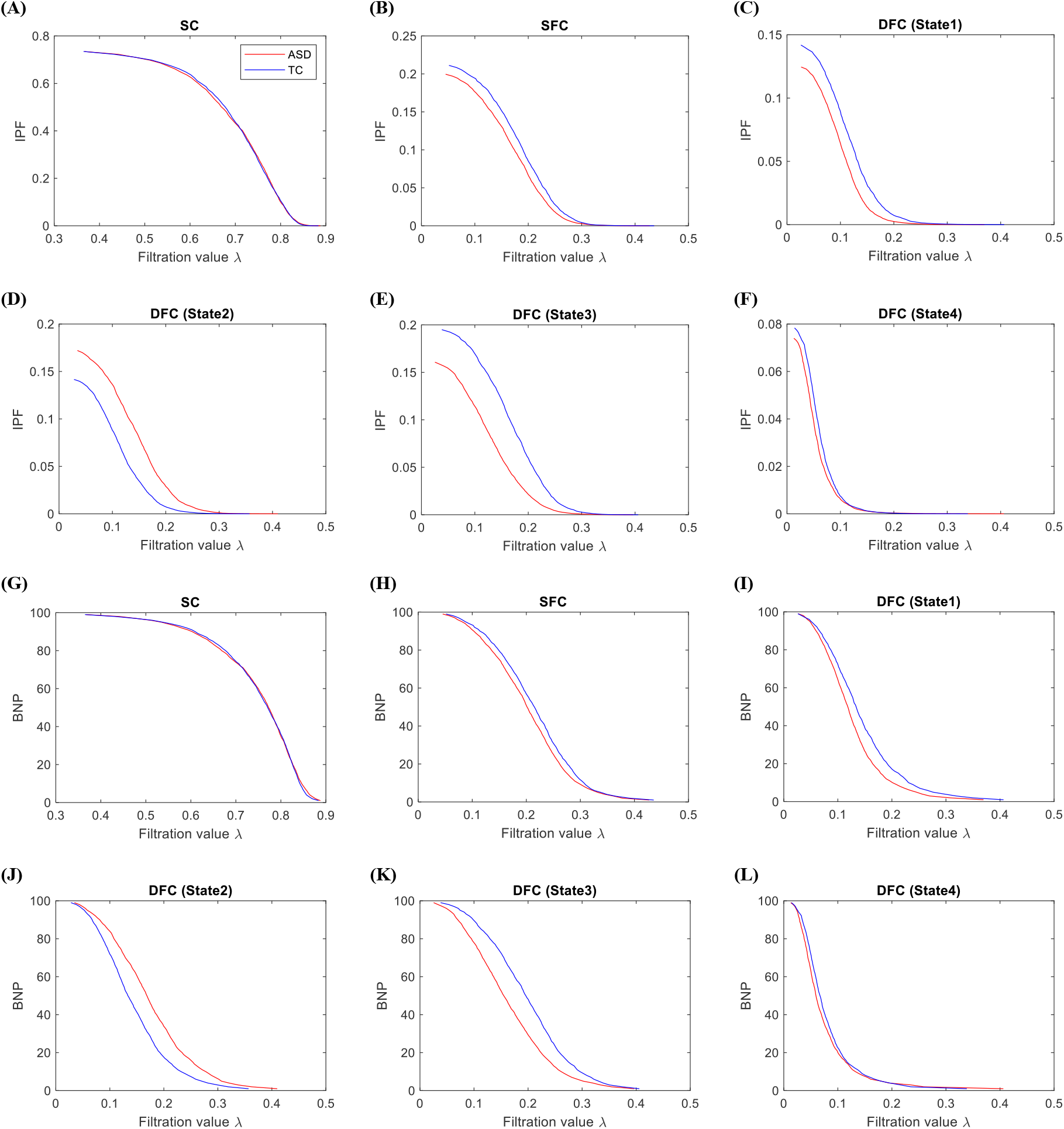
The IPF ((A)-(F)) and BNP ((G)-(L)) of ASDs and TCs overaged over subjects. Results are for SC, SFC, State1, State2, State3, and State4 of DFC. The red and blue lines represent the results of ASD and TC groups.

#### 2.6.3. PH metrics

##### Slope of integrated persistent feature (SIP)

The IPF criterion for each subject is a set of values for different values of *λ_j_*s, while for statistical analysis one wants only one value for each subject. Also, metric with one value provides a practical reference for evaluating disease progression and for effective treatments. Therefore, IPF values with respect to *λ_j_* values are fitted by a first-order linear function, and the absolute of the resulting line slope is used as the SIP metric (Kuang et al., 2019a).

##### Slope of Betti number plot (SBN)

This metric is computed like SIP. The only difference is using the BNP instead of the IPF (Lee et al., 2012). Both the SBN and SIP metrics can be thought as the information diffusion rate or convergence rate to a fully connected component (Kuang et al., 2019a).

##### IPF_AUC

This metric is the area under the curve (AUC) of the IPF criterion.

##### BNP_AUC

This metric is the area under the curve (AUC) of the BNP criterion.

##### Mean of the area of persistent landscapes for components/holes (MAPL0/ MAPL1)

If the areas enclosed below all the pl curves of a subject are added and divided by the number of pl curves, the MAPL of that subject is obtained:

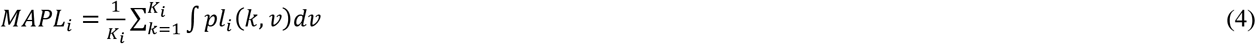

That K_i_ is the total number of pl curves of the i^th^ subject. If the pl curves of components/holes are used in (4), the value of MAPL0/MAPL1 is obtained.

##### Distance of persistent landscapes for components/holes (DPL0/DPL1)

The distance between the mean pl curve of i^th^ subject 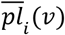 and mean pl curve of the control group 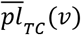 is defined as:

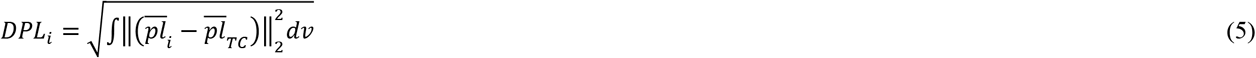

where

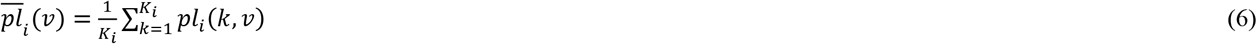

and

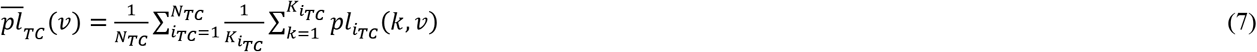

where *i_TC_* is i^th^ subject of control, and *N_TC_* is the total number of control subjects.

If the pl curves of components/holes are used in (5), the value of DPL0/DPL1 is obtained.

##### Kernel-based integrated persistent feature (KBI)

In addition to SIP, kernel-based methods are other approaches for extracting one value from the IPF curve (Kuang et al., 2019b). Kernel approaches may be a more appropriate way for computing one index value from IPF and BNP plots, as these plots are not linear (Kuang et al., 2019b). In this approach, the first, a template is computed for IPF and corresponding filtration values 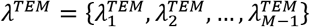 using the distance matrix averaged over subjects of the control group. The IPF values of the template are represented by *tIPF* = {*t*_1_, *t*_2_,…, *t*_*M*-1_}. If the IPF and filtration values of i^th^ subject are represented by 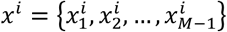 and 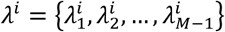 KBI is computed as

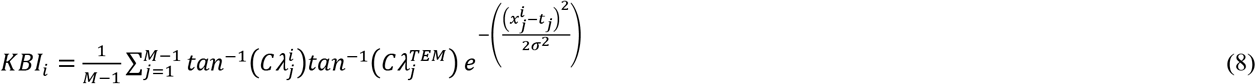

where

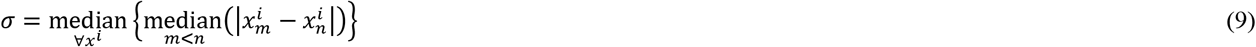

and

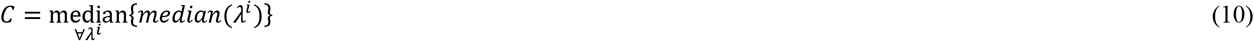

where |.| represents the absolute operator and *tan* is the tangent function.

##### Kernel-based Betti number plot (KBB)

This metric is computed like KBI. The only difference is using the BNP instead of the IPF.

### 2.7. Statistical analysis

The *t-test* was used to compare the results of two groups from a statistical point of view. Results with significant differences (*p<0.05*) were entered into the correction process. For a given metric, the non-parametric permutation testing with 1000 repetitions was performed to correct its *p-value*. In each repetition, the metric values of all studied subjects were shuffled randomly and assigned to subjects so that the number of metric values of each group was kept. The mean of the metric values was computed for each group. Then, the absolute of the subtraction between group means was computed. This process was repeated 1000 times and the computed absolute values formed a distribution. Using this distribution, the probability of the original absolute value (computed without shuffling the metric values) was computed. The probability was the ratio of the number of the absolute values which are larger than the original absolute value to 1000.

### 2.8. Classification

In this study, Support Vector Machine (SVM) (Cortes and Vapnik, 1995) with default parameters of MATLAB 2021b was implemented for classification. The classification was performed using the statistically significant features (having corrected *p* smaller than *0.05*). The results of each of the SC and SFC techniques, and each of the states of DFC was separately used for the classification if these techniques and states had significant features. Otherwise, their performance for classification was not evaluated. To evaluate the classification performance, a 5-fold cross-validation technique was performed 500 times. The values of classification metrics were averaged over 500 repetitions to obtain the final metric values. Three metrics were used, including accuracy, sensitivity, and specificity. These parameters can be calculated using the following equations:

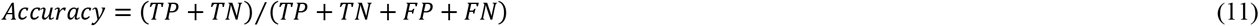

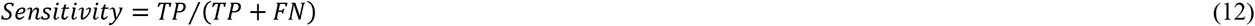

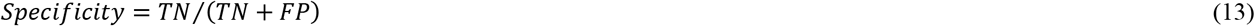

where TP, FP, TN, and FN represent the true positive, false positive, true negative, and false negative, respectively. Moreover, the receiver operating characteristic (ROC) curve was used to visually compare the performance of different techniques and states. The values of false positive rate and true positive rate are presented in the vertical and horizontal axes of the ROC diagram, respectively. In this diagram, a method with an AUC closer to 1 offers better performance.

## 3. Results

### 3.1. States of dynamic functional connectivity

The optimum cluster number for both ASD and TC groups was 4 (Figure. S1). The DFC states of ASD and TC groups are shown in Figure 4. In State1, strong positive connections can be seen for both ASD and TC groups within 7 studied networks. These connections are stronger for ASDs than TCs in the SVAN, DAN, SMN, and VN. The number and strength of connections between SVAN, DAN, and SMN are more for ASDs compared to TCs. In State2 as opposed to State1, the number and strength of within and between network connections are more for TCs when considering SVAN, DAN, SMN, and VN. The most negative links can be seen in State3. In this state, the DMN, CN, and LN have negative connections with SVAN, DAN, and SMN. Three groups of networks have positive within and between connections that exist in State3: 1) (DMN, CN, LN) 2) (SVAN, DAN, SMN) 3) VN. The number and strength of both positive and negative links are more for the ASD group. State4 is a globally hyperconnected state which has also been found in other studies (de Lacy et al., 2017; Rashid et al., 2018; Mash et al., 2019) and may be predominated by partially nonneuronal global signal fluctuations (Mash et al., 2019). Global signal has been shown to contain meaningful neural information (Schölvinck, Maier, Ye, Duyn, and Leopold, 2010). Thus, it has not been removed during preprocessing and may justify the appearance of state4.

**Figure 4.**
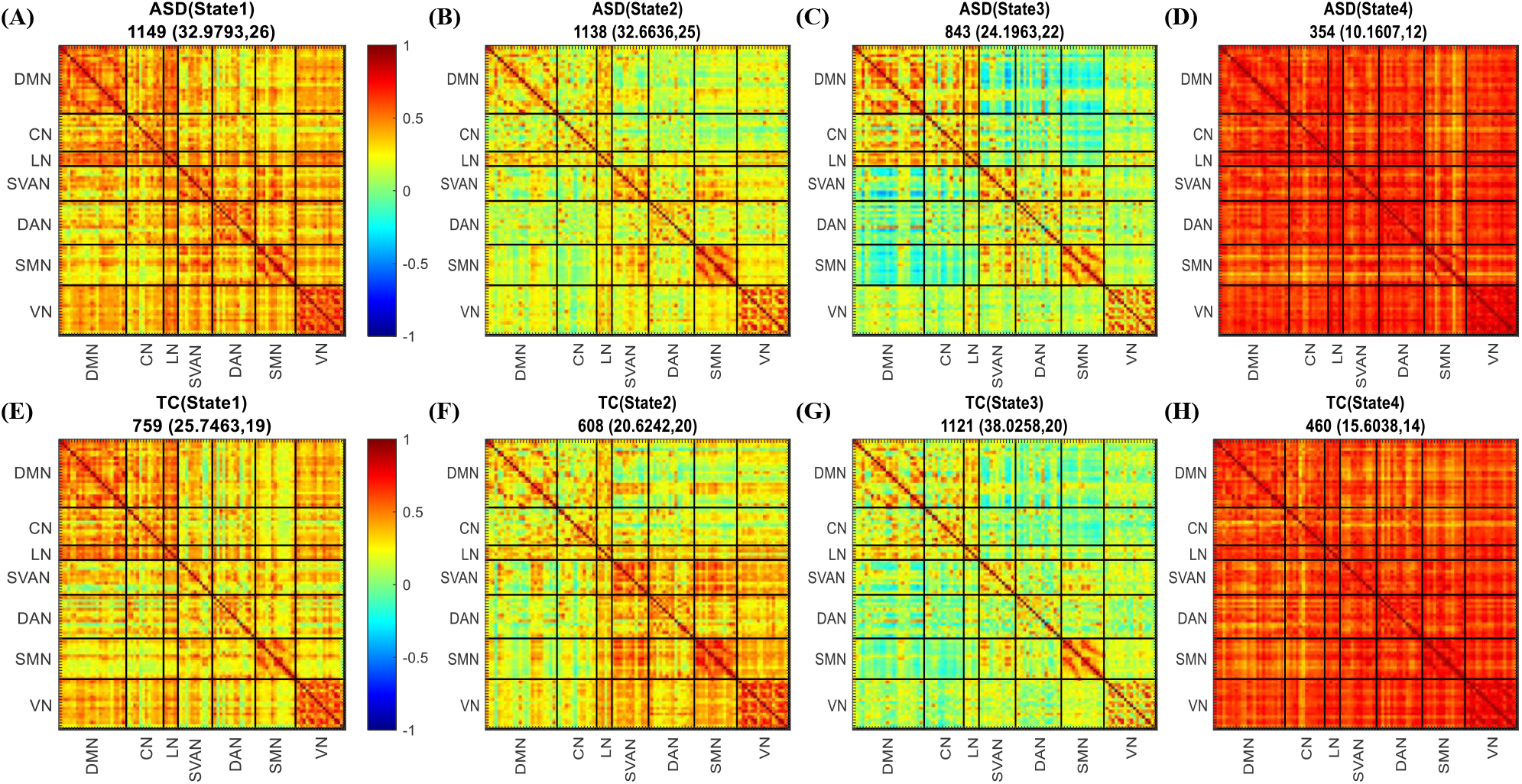
Four states of ASD (first row) and TC (second row) groups. There are numbers above each state written in the format of **A**(**B**, **C**): **A** is the number of FC matrices contributed to the given state, **B** represents the ratio of **A** to the total number of FC matrices of the studied group (in percent), **C** is the number of subjects contributed in the given state.

### 3.2. Results of graph metrics

The results of the statistical comparison are listed in Table 2. The statistically significant differences between ASDs and TCs are only seen in the States1-3 of DFC.

**Table 2.** The statistical results when comparing segregation and integration metrics between ASDs and TCs. The t of t-test and corrected *p-values* are reported. The significant *p-values* (<0.05) are written in bold style.

|  | SC | SFC | State1 | State2 | State3 | State4 |
| --- | --- | --- | --- | --- | --- | --- |
|  | *t* (*p*) | *t* (*p*) | *t* (*p*) | *t* (*p*) | *t* (*p*) | *t* (*p*) |
| **GE** | 0.35 (0.35) | 0.26 (0.41) | 2.1 (**0.02**) | -4 (**<3e-4**) | 3 (**0.004**) | 0.5 (0.29) |
| **ME** | 0.3 (0.36) | -0.25 (0.39) | -2.84 (**0.003**) | -0.26 (0.8) | 2.2 (**0.02**) | 0.38 (0.37) |
| **Radius** | 0.21 (0.37) | -0.44 (0.3) | -3.18 (**<e-3**) | 0.77 (0.44) | 0.85 (0.4) | 0.55 (0.32) |
| **Diameter** | 0.38 (0.34) | -0.29 (0.38) | -1.58 (0.12) | -1.32 (0.19) | 2.83 (**0.003**) | -0.03 (0.45) |
| **AC** | 0.18 (0.43) | 0.1 (0.46) | -0.28 (0.78) | -0.57 (0.57) | 0.5 (0.6) | -1.04 (0.14) |
| **MCC** | -0.39 (0.34) | -0.18 (0.43) | -4.06 (**<2e-4**) | 4.97 (**<2e-5**) | -1.64 (0.1) | -0.04 (0.48) |
| **MEC** | 0.4 (0.35) | 0.67 (0.25) | 0.8 (0.4) | -1.32 (0.19) | -0.76 (0.44) | 0.32 (0.38) |
| **Modularity** | 0.3 (0.36) | -0.5 (0.32) | 1.5 (0.13) | 0.53 (0.6) | 0.26 (0.79) | 0.007 (0.48) |
| **MS** | -0.1 (0.4) | 0.15 (0.44) | 3.85 (**<4e-4**) | -4.6 (**<4e-5**) | 0.11 (0.9) | -0.06 (0.48) |

Abbreviations: GE = global efficiency; ME = mean of eccentricity; AC = Assortativity coefficient; MCC = mean of clustering coefficient; MEC = mean of eigenvector centrality; MS = mean of strength; SC = structural connectivity; SFC = static functional connectivity.

Significant metrics in State2 show contrary behavior to their corresponding metrics in State1 when comparing ASD with TC. This means that the two metrics GE and MS are significantly less/more for ASDs than TCs in the State2/State1 with statistical information of (all *p<3e-4*, all *t<-4*)/(all *p<0.02*, all *t>2.1*). In contrast, MCC is significantly more/less for ASDs than TCs in the State2/State1 with statistical information of (*p=2e-5, t=4.97*)/(*p=2e-4, t=-4.06*).Also, in State1, the ME and Radius were significantly less for ASDs than TCs (all *p<0. 003*, all *t<-2.84*). In State3, the GE, ME, and Diameter metrics are significantly more for ASDs compared to TCs (all *p<0.02*, all *t>2.2*).

### 3.3. Results of persistent homology metrics

The results of the statistical comparison are listed in Table 3. Most of the statistically significant differences between ASDs and TCs are seen in the States1-3 of DFC.

**Table 3.** The statistical results when comparing persistent homology metrics between ASDs and TCs. The t of t-test and corrected *p-values* are reported. The significant *p-values* (*<0.05*) are written in bold style.

|  | SC | SFC | State1 | State2 | State3 | State4 |
| --- | --- | --- | --- | --- | --- | --- |
| | $t$ ($p$) | $t$ ($p$) | $t$ ($p$) | $t$ ($p$) | $t$ ($p$) | $t$ ($p$) |
| SIP | -0.59 (0.59) | -1.42 (0.16) | -0.22 (0.82) | 1.32 (0.19) | -3.76 (**<6e-4**) | -1.62 (0.11) |
| IPF_AUC | -0.26 (0.79) | -1.25 (0.22) | -2.3 (**0.007**) | 3.52 (**0.001**) | -4.03 (**<3e-4**) | -1.07 (0.29) |
| SBN | -0.68 (0.49) | -0.1 (0.91) | 2 (**0.02**) | -2.55 (**0.004**) | -0.2 (0.83) | -1.48 (0.15) |
| BNP_AUC | -0.13 (0.89) | -0.57 (0.57) | -2.86 (**0.002**) | 3.77 (**<5e-5**) | -2.89 (**<e-3**) | -0.58 (0.56) |
| MAPL0 | -0.61 (0.54) | -0.94 (0.35) | -3.72 (**<6e-4**) | 3.08 (**0.003**) | -0.94 (0.35) | 0.28 (0.77) |
| MAPL1 | 1.41 (0.16) | -0.62 (0.53) | -2.78 (**0.007**) | 1.98 (0.054) | -0.15 (0.88) | 0.32 (0.75) |
| DPL0 | 0.23 (0.82) | -1.08 (0.28) | 2.16 (**0.01**) | 1.35 (0.18) | -0.86 (0.39) | 0.32 (0.75) |
| DPL1 | 0.86 (0.39) | -1.26 (0.21) | -1.05 (0.29) | 2.38 (**0.01**) | 0.44 (0.66) | 0.81 (0.42) |
| KBI | -1.85 (0.07) | 0.7 (0.48) | -2.75 (**0.003**) | 0.76 (0.44) | -0.76 (0.44) | 0.44 (0.66) |
| KBB | -2.12 (**0.02**) | 1.75 (0.08) | -1.22 (0.23) | -3.33 (**0.002**) | 3.02 (**0.003**) | 1.87 (0.07) |

Abbreviations: SIP = slope of integrated persistent feature; IPF\_AUC = area under curve of integrated persistent feature; SBN = slope of betti number plot; BNP\_AUC = area under curve of betti number plot; MAPL = mean area of persistent landscapes; DPL = distance of persistent landscapes; KBI = kernel-based integrated persistent feature; KBB = kernel-based betti number plot.

Most of the significant metrics in State2 offer larger values for ASDs than TCs whereas these metrics in State1 offer smaller values for ASDs compared to TCs. These metrics are IPF_AUC, BNP_AUC, and MAPL0 with (all *p<0.01*, all *t>2.38*)/(*all p<0.007*, all *t<-2.3*) in State2/State1. In State1, significant results can be also seen for MAPL1, DPL0, SBN, and KBI with (*p=0.007, t=-2.78*), (*p=0.01, t=2.16*), (*p=0.02, t=2*), and (*p=0.003, t=-2.75*), respectively. These results show that most of the PH metrics in State1 are significantly less for ASDs than TCs. In State2, SBN, DPL1, and KBB are significantly different between ASD and TC groups with (*p=0.004, t=-2.55*), (*p=0.01, t=2.38*), and (*p=0.002, t=-3.33*), respectively. Overall, States 1 and 2 are opposite to each other in terms of most of the PH metrics. In State3, significant results are for metrics related to IPF and BNP values. These metrics are SIP (*p=6e-4, t=-3.76*), IPF_AUC (*p=3e-4, t=-4.03*), BNP_AUC (*p=e-3, t=-2.89*), and KBB (*p=0.003, t=3.02*). On average, the ASDs compared to TCs have lower SIP values and, consequently, lower IPF_AUC values in this state.

The IPF and BNP plots (Figure 3) also show that the differences between ASD and TC groups in States1-3 are more than State4, SFC, and SC.

### 3.4. Results of classification

The States1-3 provided significant results for both graph and PH metrics. Hence, the classification was performed only for these states and using their corresponding significant metrics. The results obtained using the SVM classifier for graph and PH metrics are summarized in Table 4.

**Table 4.** The classification results when classifying ASDs and TCs using linear SVM. Classification is performed only in State1, State2, and State3 of DFC due to offering statistically significant features. For each state, only the significant features (p<0.05) are employed for classification.

|  |  | State1 | State2 | State3 |
| --- | --- | --- | --- | --- |
| Graph metrics | Accuracy (%) | 73.33 | 71.11 | 73.8 |
|  | Sensitivity (%) | 57.89 | 55 | 70 |
|  | Specificity (%) | 84.61 | 84 | 77.27 |
| PH metrics | Accuracy (%) | 71.11 | **100** | **100** |
|  | Sensitivity (%) | 57.89 | **100** | **100** |
|  | Specificity (%) | 80.77 | **100** | **100** |

PH = persistent homology.

The PH metrics in States2-3 offer the best and highest performance i.e., all three metrics, including Accuracy, Specificity, and Sensitivity are 100%. Based on these results statistical results, it can be said that States2-3 compared to States1,4, SC, and SFC are more informative when the purpose is discriminating between ASD and TC subjects. Also, based on the classification results, it can be said that PH metrics are more informative than graph global metrics.

In addition to quantitative parameters, the performance of graph and PH metrics has been investigated using the ROC curve. As shown in Figure 5, the highest performances (AUC=1) are seen for PH metrics in States 2 and 3. Based on the AUC values, it can be said that State2 is the only state offering the best performance for both graph and PH.

**Figure 5.**
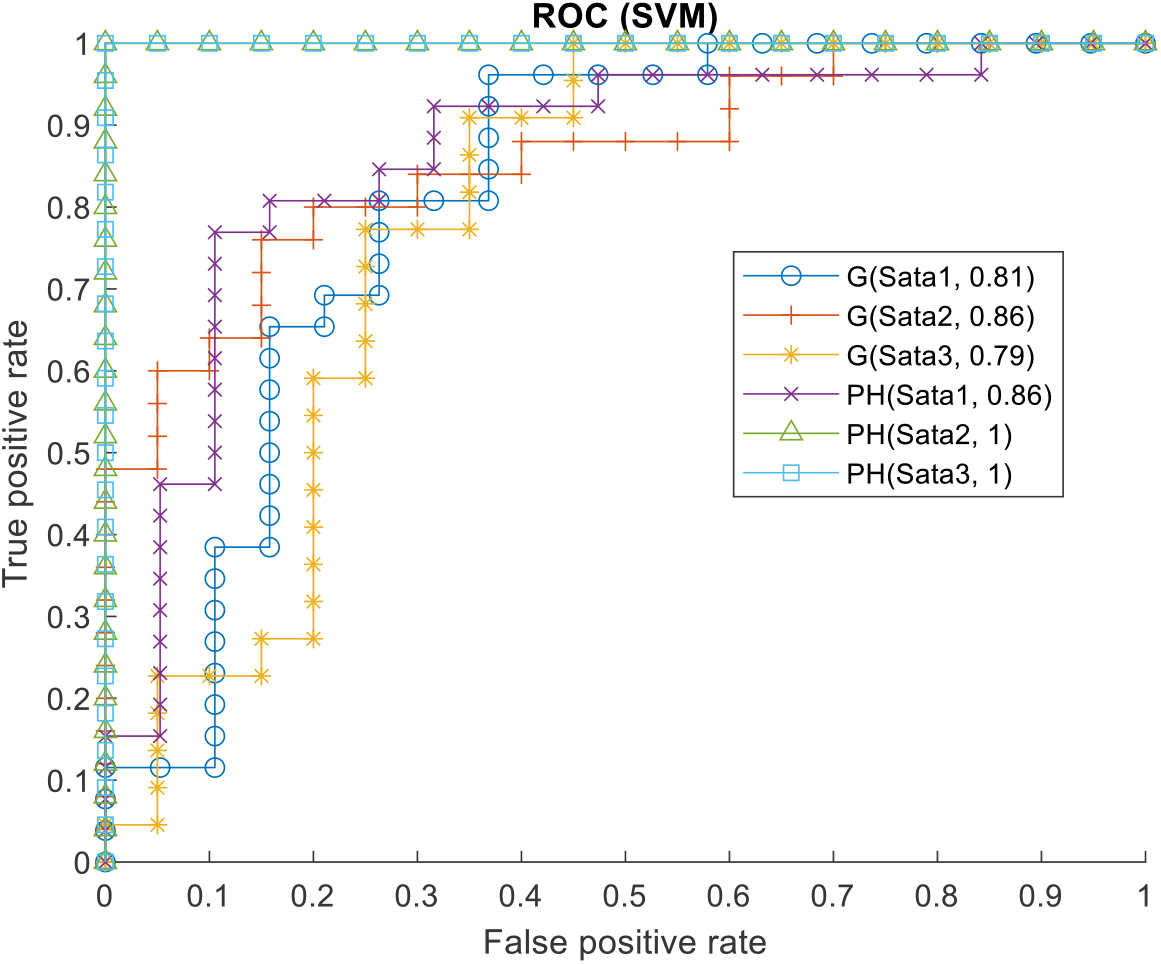
Comparisons of ROC curves. The AUC values are written inside the legend. The persistent homology (PH) metrics offer the best performance in State2 and State3 of DFC. The character G stands for graph metrics.

## 4. Discussion

In this study, some of the graph global metrics were used to investigate the segregation and integration of brain networks. Also, some metrics of PH were defined using the components and holes of the network. The PH metrics allow addressing dynamic abnormalities at a more global level, where the evolution of the global pattern of connectivity contributed by all scales (filtration values) is probed. These metrics were applied to SC, SFC, and states of DFC, separately. For evaluations, statistical analysis and classification were performed. Results showed that only three states (States 1-3) of DFC offered statistically significant features. This finding may show that DFC can be more informative than SC and SFC from a topological point of view. The classification results in States 2 and 3 showed that PH metrics could provide higher performance than graph metrics for discrimination of ASDs and TCs. These results highlighted the key roles of States 2 and 3 and the advantages of PH metrics.

The k-means clustering technique offered four states for each of the ASD and TC groups, which are shown in Figure 4. In State1, the ASD group had much more positive links with higher strength. These links were both within and between networks, including SVAN, DAN, SMN, and VN. State2 showed a contrary behavior to States1. In State2, the TC group compared to the ASD group had much more positive links with higher strength. These differences were also seen within and between SVAN, DAN, SMN, and VN. Such opposite patterns of State1 and State2 showed themselves in the studied metrics, where many of the graph and PH metrics offered significant opposite results in these states. The State2 showed isolation of SVAN, DAN, SMN, and VN for ASDs. In State3, three modules of positive links could be seen, including Module1: ROIs of DMN, CN, and LN; Module2: ROIs of SVAN, DAN, and SMN; Module3: ROIs of VN. Negative connections existed between Module1 and Module2. In this state, the strength of both positive and negative links in ASDs was more than TCs. As a result, the modular structure was more apparent for ASDs compared to TCs. State4 was a hyperconnectivity state. This state may be for nonneuronal global signal fluctuations (Mash et al., 2019). There were seen no significant results in this state.

In State1, ASDs had significantly lower ME (*p=0.003, t=-2.84*). This meant that ROIs of ASDs were closer to each other than TCs (more integration). The radius results also confirmed this idea (*p<e-3, t=-3.18*). In this state, the number and strength of positive links of ASDs were more than TCs which in turn resulted in such closeness (Figure 4). Such differences in positive links existed in both within and between network connections of SVAN, DAN, SMN, and VN. In State3, contrary behavior was seen, where having more negative links with higher strengths resulted in significantly higher ME (*p=0.02, t=2.2*) and Diameter (*p=0.003, t=2.83*) for ASDs compared to TCs. This meant that the ROIs of ASDs were segregated more than TCs in this state. These negative links and segregation were between (DMN, CN, LN) and (SVAN, DAN, SMN).

The GE results of Table 2 demonstrated that ASDs exchange information more/less efficiently than TCs in State1/State2. This contrary behavior of efficiency metric can be justified by considering the images of these states shown in Figure 4. The ASD/TC group had more positive/more positive and less negative connections in State1/State2 than TC/ASD group. In State 3, it can be said that the positive links of this state compensated for the effect of negative links and improved the efficiency of ASDs. As a result, the GE was significantly more for ASDs in State3.

The average clustering coefficient of a network is a direct metric for measuring the local specialization of the network (Bullmore and Sporns, 2009). Thus, the results of the clustering coefficient in State2 may show a decrease in brain functional specialization in ASDs due to having more local connections rather than global ones. Similar results have also been reported in the electroencephalography (EEG) study in the theta, beta, and low gamma bands (Ye et al., 2014). The results of the strength metric showed the over-connectivity/under-connectivity in ASDs compared to TCs in the State1/State2 (Ye et al., 2014; Supekar et al., 2013; Hull et al., 2016; Di Martino et al., 2013).

The SBN of ASD/TC was significantly steeper than TC/ASD in State1/State2 with statistical information (*p=0.02, t=2)/(p=0.004, t=-2.55*). These results may show that the information diffusion speed in the ASD group is higher/lower than normal in the State1/State2 (Kuang et al., 2019a). SBN is directly related to the whole-brain topology. As a result, it can be said that the SBN results speak about global topological abnormalities of ASDs in States 1 and 2. Therefore, it can be speculated that the higher/lower diffusion speed is a result of the increased/decreased functional integration of the whole brain, which may further lead to cognitive defects in patients with ASD. The connectivity maps of State1 and State2 (Figure4) demonstrated these increased and decreased integration in State1 and State2, respectively. The ME, Radius, and MS metrics of the graph confirm this speculation in State1, where ME and MS were significantly lower and higher for ASDs compared to TCs, respectively. The connectivity map of State2 (Figure 4) also confirms the higher integration of TCs (higher isolation of ASDs). SIP like as SBN indicates the information diffusion rate. In State3, the SIP of ASDs was significantly less than TCs (*p<6e-4, t=-3.76*). This may be due to having many more negative links (increased segregation). The results of ME and Diameter are in line with the SIP result in this state.

The positive/negative statistical value for MAPL and DPL metrics means that the number and persistency of components and holes are more/less for ASDs than TCs. Therefore, global topological abnormalities can be concluded for ASDs.

The best performance of classification was obtained using PH metrics in States 2 and 3. The IPF_AUC, BNP_AUC, and KBB metrics of PH were significant in both States 2 and 3. Therefore, it may be said that these metrics offer the most important discriminative features when classifying ASDs with TCs. These three metrics had contrary behavior in States 2 and 3. Also, in these two states, the segregation and isolation of ASDs were more than TCs. Also, the results of SBN and SIP suggested that the data diffusion rate of ASDs compared to TCs was lower in these states. Totally, it may be concluded that IPF and BNP based metrics play important role in discriminating ASDs from TCs in dynamic states where the networks of ASDs compared to controls are more segregated and isolated.

All the significant results of both graph and PH metrics provide some empirical evidence for the impaired global organization and disrupted neuronal integration in ASD.

Overall, the DFC analysis provides more alterations of functional connectivity, which cannot be revealed by SFC analysis. Findings of this study suggest that the brain DFC network analysis based on persistent homology is more likely to distinguish ASD from TC, and is more likely to explore potential biomarkers for ASD imaging. Our current results should be viewed as exploratory and need to be further confirmed in other independent cohorts in future.

### 4.1. Limitations and future directions

One of the limitations of this work was studying only male subjects. Thus, the effect of gender was not investigated on the studied measures. For future works, the measures of this paper can be applied to the rfMRI and DTI data of both males and females. Using these metrics for task fMRI data of ASD and TC can be another topic for future works.

In this study, the classification was performed to reveal the capacity of graph and PH metrics and different states of DFC in discriminating of ASDs from TCs. The value 1 of specificity and sensitivity of Table 4 does not mean that PH metrics in State2 and State3 always offer 1 for mentioned metrics. It just shows that State2 and State3 are more informative than the two rest states. Also, it demonstrates that PH metrics can be more valuable than graph metrics under the same conditions. Moreover, it suggests that it is worthwhile to devote some of the future research to studying the PH metrics in the states of DFC. In future work, one can use a much larger dataset to report much more reliable and accurate values for classification metrics.

## 5. Conclusion

In this study, the ability of SC, SFC, and states of DFC in discriminating between ASDs and TCs was investigated using the global metrics of graph and metrics of PH. For PH, the components and holes of the distance matrix were studied. The studied metrics investigated brain function and structure from a topological point of view. For the first time, these metrics were used to study the states of DFC of autism subjects, consequently, a novel insight into the whole-brain network analysis was offered in this study. Four states were found for ASDs and TCs when using DFC analysis. Significant discriminative features were only found using DFC analysis and in the States1-3. The best classification performance was seen in States 2 and 3. The classification results showed that PH metrics had significantly higher discriminative capacity than graph metrics in States 2 and 3. The contrary behavior was seen between State1 and State2 in terms of both graph and PH significant metrics. All these results demonstrated the promising perspective of using the PH approach and DFC analysis for attaining discriminative features and potential imaging biomarkers when studying ASDs in comparison to TCs.

## Availability of data and codes

The datasets and codes used during the current study are available from the corresponding author upon request.

## Approval for human experiments

For the included data repositories SDSU, the following Institutional Review Boards (IRB) and ethics committee have approved the experiments, and data acquisition procedures were following their guidelines and regulations, respectively: Institutional Review Board Operations at San Diego State University’s Human Research Protection Program (HRPP). Furthermore, all experiments were following HIPAA guidelines and 1000 Functional Connectomics Project/INDI protocols, that is, all data were fully anonymized with no protected health information and legal guardians of all participants have signed informed consent.

## Supplementary material

Supplementary material is available at *CIBM* online.

## References

Adams, H., Tausz, A., & Vejdemo-Johansson, M. 2014. JavaPlex: A research software package for persistent (co) homology. In International congress on mathematical software (pp. 129–136). Springer, Berlin, Heidelberg.

Aggarwal, P., Gupta, A., 2019. Multivariate graph learning for detecting aberrant connectivity of dynamic brain networks in autism. Medical image analysis, 56, 11–25.

Allen, E. A., Damaraju, E., Plis, S. M., Erhardt, E. B., Eichele, T., Calhoun, V. D., 2014. Tracking whole-brain connectivity 33 dynamics in the resting state. Cerebral Cortex, 24(3), 663–676.

American Psychiatric Association, 2013. Diagnostic and statistical manual of mental disorders. American Psychiatric Association, Washington DC.

Arthur, D., & Vassilvitskii, S. 2006. k-means++: The advantages of careful seeding. Stanford.

Biswal, B.B., 2012. Resting state fMRI: a personal history. Neuroimage 62, 938–944.

Bullmore E., Sporns O. 2009. Complex brain networks: graph theoretical analysis of structural and functional systems. Nat. Rev. Neurosci. 10, 186–198. 10.1038/nrn2575.

Bubenik, P. 2015. Statistical topological data analysis using persistence landscapes. J. Mach. Learn. Res., 16(1), 77–102.

Caballero-Gaudes, C., Reynolds, R. C., 2017. Methods for cleaning the BOLD fMRI signal. Neuroimage, 154, 128–149.

Carlsson, G. 2009. Topology and data. Bulletin of the American Mathematical Society, 46(2), 255–308.

Cheng, W., Rolls, E. T., Gu, H., Zhang, J., Feng, J., 2015. Autism: reduced connectivity between cortical areas involved in face expression, theory of mind, and the sense of self. Brain, 138(5), 1382–1393.

Choi, H., Kim, Y. K., Kang, H., Lee, H., Im, H. J., Kim, E. E.,… & Lee, D. S. 2014. Abnormal metabolic connectivity in the pilocarpine-induced epilepsy rat model: a multiscale network analysis based on persistent homology. Neuroimage, 99, 226–236.

Chung, M. K., Hanson, J. L., Ye, J., Davidson, R. J., & Pollak, S. D. 2015. Persistent homology in sparse regression and its application to brain morphometry. IEEE transactions on medical imaging, 34(9), 1928–1939.

Cortes, C., & Vapnik, V. 1995. Support-vector networks. Machine Learning, 20(3), 273–297.

De Lacy, N., Doherty, D., King, B. H., Rachakonda, S., & Calhoun, V. D. 2017. Disruption to control network function correlates with altered dynamic connectivity in the wider autism spectrum. NeuroImage: Clinical, 15, 513–524.

Di Martino, A., Zuo, X. N., Kelly, C., Grzadzinski, R., Mennes, M., Schvarcz, A., Milham, M. P. 2013. Shared and distinct intrinsic functional network centrality in autism and attention-deficit/hyperactivity disorder. Biological psychiatry, 74(8), 623–632.

De Vico Fallani F., Richiardi J., Chavez M., Achard S. 2014. Graph analysis of functional brain networks: practical issues in translational neuroscience. Philos. Trans. R. Soc. Lond. B Biol. Sci. 369:20130521. 10.1098/rstb.2013.0521

Drysdale, A. T., Grosenick, L., Downar, J., Dunlop, K., Mansouri, F., Meng, Y., Schatzberg, A. F., 2017. Resting-state connectivity biomarkers define neurophysiological subtypes of depression. Nature medicine, 23(1), 28–38.

Edelsbrunner, H., Harer, J. L. 2010. Computational topology: an introduction. American Mathematical Society: Heidelberg, Germany.

Elsabbagh, M., Divan, G., Koh, Y. J., Kim, Y. S., Kauchali, S., Marcín, C., Fombonne, E. 2012. Global prevalence of autism and other pervasive developmental disorders. Autism research, 5(3), 160–179.

Farahani, F. V., Karwowski, W., & Lighthall, N. R. 2019. Application of graph theory for identifying connectivity patterns in human brain networks: a systematic review. frontiers in Neuroscience, 13, 585.

Fornito, A., Zalesky, A., Bullmore, E. 2016. Fundamentals of brain network analysis. Academic Press.

Fu, Z., Tu, Y., Di, X., Du, Y., Sui, J., Biswal, B. B.,… Calhoun, V. D. 2018. Transient increased thalamic-sensory connectivity and decreased whole-brain dynamism in autism. Neuroimage. https://doi.org/10.1016/j.neuroimage.2018.06.003.

Goldani, A. A., Downs, S. R., Widjaja, F., Lawton, B., & Hendren, R. L. 2014. Biomarkers in autism. Frontiers in psychiatry, 5, 100.

Grecucci, A., Siugzdaite, R., Job, R., 2017. Advanced Neuroimaging Methods for Studying Autism Disorder. Frontiers in neuroscience, 11, 533.

Ha, S., Sohn, I. J., Kim, N., Sim, H. J., Cheon, K. A., 2015. Characteristics of brains in autism spectrum disorder: structure, function and connectivity across the lifespan. Experimental neurobiology, 24(4), 273–284.

Hull J. V., Dokovna L. B., Jacokes Z. J., Torgerson C. M., Irimia A., Van Horn J. D. 2016. Resting-state functional connectivity in autism spectrum disorders: a review. Front. Psychiatry 7:205. 10.3389/fpsyt.2016.00205.

Hull, J. V., Dokovna, L. B., Jacokes, Z. J., Torgerson, C. M., Irimia, A., Van Horn, J. D., 2017. Resting-state functional connectivity in autism spectrum disorders: A review. Frontiers in psychiatry, 7, 205.

Iidaka, T., 2015. Resting state functional magnetic resonance imaging and neural network classified autism and control. Cortex, 63, 55–67.

Itahashi, T., Yamada, T., Watanabe, H., Nakamura, M., Jimbo, D., Shioda, S., Hashimoto, R. 2014. Altered network topologies and hub organization in adults with autism: a resting-state fMRI study. PloS one, 9(4), e94115.

Jou, R. J., Jackowski, A. P., Papademetris, X., Rajeevan, N., Staib, L. H., Volkmar, F. R., 2011. Diffusion tensor imaging in autism spectrum disorders: preliminary evidence of abnormal neural connectivity. Australian New Zealand Journal of Psychiatry, 45(2), 153–162.

Keown, C. L., Datko, M. C., Chen, C. P., Maximo, J. O., Jahedi, A., & Müller, R. A. 2017. Network organization is globally atypical in autism: a graph theory study of intrinsic functional connectivity. Biological Psychiatry: Cognitive Neuroscience and Neuroimaging, 2(1), 66–75.

Kuang, L., Han, X., Chen, K., Caselli, R. J., Reiman, E. M., Wang, Y., & Alzheimer’s Disease Neuroimaging Initiative. 2019a. A concise and persistent feature to study brain resting-state network dynamics: Findings from the Alzheimer’s Disease Neuroimaging Initiative. Human brain mapping, 40(4), 1062–1081.

Kuang, L., Zhao, D., Xing, J., Chen, Z., Xiong, F., & Han, X. 2019b. Metabolic brain network analysis of FDG-PET in Alzheimer’s disease using kernel-based persistent features. Molecules, 24(12), 2301.

Kuang, L., Gao, Y., Chen, Z., Xing, J., Xiong, F., & Han, X. 2020a. White matter brain network research in Alzheimer’s disease using persistent features. Molecules, 25(11), 2472.

Kuang, L., Jia, J., Zhao, D., Xiong, F., Han, X., Wang, Y., & Alzheimer’s Disease Neuroimaging Initiative. 2020. Default mode network analysis of APOE genotype in cognitively unimpaired subjects based on persistent homology. Frontiers in Aging Neuroscience, 12, 188.

Lee, H., Kang, H., Chung, M. K., Kim, B. N., & Lee, D. S. 2012. Persistent brain network homology from the perspective of dendrogram. IEEE transactions on medical imaging, 31(12), 2267–2277.

Lee, H., Kang, H., Chung, M. K., Lim, S., Kim, B. N., Lee, D. S. 2017. Integrated multimodal network approach to PET and MRI based on multidimensional persistent homology. Human Brain Mapping, 38(3), 1387–1402.

Li, D., Karnath, H. O., Xu, X., 2017. Candidate biomarkers in children with autism spectrum disorder: a review of MRI studies. Neuroscience bulletin, 33(2), 219–237.

Mash, L. E., Reiter, M. A., Linke, A. C., Townsend, J., Müller, R. A., 2018. Multimodal approaches to functional connectivity in autism spectrum disorders: an integrative perspective. Developmental neurobiology, 78(5), 456–473.

Mash, L. E., Linke, A. C., Olson, L. A., Fishman, I., Liu, T. T., Müller, R. A., 2019. Transient states of network connectivity are atypical in autism: A dynamic functional connectivity study. Human brain mapping, 40(8), 2377–2389.

Mukherjee, P., McKinstry, R. C., 2006. Diffusion tensor imaging and tractography of human brain development. Neuroimaging Clinics, 16(1), 19–43.

Nagae, L. M., Zarnow, D. M., Blaskey, L., Dell, J., Khan, S. Y., Qasmieh, S., Roberts, T. P. L., 2012. Elevated mean diffusivity in the left hemisphere superior longitudinal fasciculus in autism spectrum disorders increases with more profound language impairment. American Journal of Neuroradiology, 33(9), 1720–1725.

Nair, A., Treiber, J. M., Shukla, D. K., Shih, P., Müller, R. A., 2013. Impaired thalamocortical connectivity in autism spectrum disorder: a study of functional and anatomical connectivity. Brain, 136(6), 1942–1955.

Otter, N., Porter, M. A., Tillmann, U., Grindrod, P., & Harrington, H. A. 2017. A roadmap for the computation of persistent homology. EPJ Data Science, 6, 1–38.

Power, J. D., Barnes, K. A., Snyder, A. Z., Schlaggar, B. L., Petersen, S. E., 2012. Spurious but systematic correlations in functional connectivity MRI networks arise from subject motion. Neuroimage, 59(3), 2142–2154.

Preti, M. G., Bolton, T. A., Van De Ville, D., 2017. The dynamic functional connectome: State-of-the-art and perspectives. NeuroImage, 160, 41–54. https://doi.org/10.1016/j.neuroimage.2016.12.061.

Rashid, B., Blanken, L. M., Muetzel, R. L., Miller, R., Damaraju, E., Arbabshirani, M. R., Tiemeier, H., 2018. Connectivity dynamics in typical development and its relationship to autistic traits and autism spectrum disorder. Human brain mapping, 39(8), 3127–3142.

Redcay, E., Moran, J. M., Mavros, P. L., Tager-Flusberg, H., Gabrieli, J. D., & Whitfield-Gabrieli, S. 2013. Intrinsic functional network organization in high-functioning adolescents with autism spectrum disorder. Frontiers in human neuroscience, 7, 573.

Rubinov, M., & Sporns, O. 2010. Complex network measures of brain connectivity uses and interpretations. Neuroimage, 52(3), 1059–1069.

Rudie, J. D., Brown, J. A., Beck-Pancer, D., Hernandez, L. M., Dennis, E. L., Thompson, P. M., Dapretto, M. J. N. C. 2013. Altered functional and structural brain network organization in autism. NeuroImage: clinical, 2, 79–94.

Song, Y., Epalle, T. M., & Lu, H. 2019. Characterizing and predicting autism spectrum disorder by performing resting-state functional network community pattern analysis. Frontiers in human neuroscience, 13, 203.

Schaefer, A., Kong, R., Gordon, E. M., Laumann, T. O., Zuo, X. N., Holmes, A. J., & Yeo, B. T., 2018. Local-global parcellation of the human cerebral cortex from intrinsic functional connectivity MRI. Cerebral cortex, 28(9), 3095–3114.

Schölvinck, M. L., Maier, A., Frank, Q. Y., Duyn, J. H., & Leopold, D. A. 2010. Neural basis of global resting-state fMRI activity. Proceedings of the National Academy of Sciences, 107(22), 10238–10243.

Sundaram, S. K., Kumar, A., Makki, M. I., Behen, M. E., Chugani, H. T., Chugani, D. C., 2008. Diffusion tensor imaging of frontal lobe in autism spectrum disorder. Cerebral cortex, 18(11), 2659–2665.

Supekar, K., Uddin, L. Q., Khouzam, A., Phillips, J., Gaillard, W. D., Kenworthy, L. E., Menon, V. 2013. Brain hyperconnectivity in children with autism and its links to social deficits. Cell reports, 5(3), 738–747.

Tax, C. M., Otte, W. M., Viergever, M. A., Dijkhuizen, R. M., & Leemans, A., 2015. REKINDLE: robust extraction of kurtosis INDices with linear estimation. Magnetic resonance in medicine, 73(2), 794–808.

Uddin, L. Q., Supekar, K., Menon, V., 2013. Reconceptualizing functional brain connectivity in autism from a developmental perspective. Frontiers in.

Watanabe, T., & Rees, G. 2017. Brain network dynamics in highfunctioning individuals with autism. Nature Communications, 8, 16048. https://doi.org/10.1038/ncomms16048.

Woo, C. W., Wager, T. D., 2015. Neuroimaging-based biomarker discovery and validation. Pain, 156(8), 1379.

Xing, J., Jia, J., Wu, X., & Kuang, L. 2022. A Spatiotemporal Brain Network Analysis of Alzheimer’s Disease Based on Persistent Homology. Frontiers in Aging Neuroscience, 14.

Yeo, B. T., Krienen, F. M., Sepulcre, J., Sabuncu, M. R., Lashkari, D., Hollinshead, M. & Buckner, R. L., 2011. The organization of the human cerebral cortex estimated by intrinsic functional connectivity. Journal of neurophysiology.

Ye, A. X., Leung, R. C., Schäfer, C. B., Taylor, M. J., & Doesburg, S. M. 2014. Atypical resting synchrony in autism spectrum disorder. Human brain mapping, 35(12), 6049–6066.

Yoo, K., Lee, P., Chung, M. K., Sohn, W. S., Chung, S. J., Na, D. L., Jeong, Y. 2017. Degree-based statistic and center persistency for brain connectivity analysis. Human Brain Mapping, 38(1), 165–181.

Zomorodian, A., & Carlsson, G. 2005. Computing persistent homology. Discrete & Computational Geometry, 33(2), 249–274.

