## Supplementary File for "Topological analysis of brain dynamics in autism based on graph and persistent homology"

### S.1. Elbow criterion for k-means clustering of LVFS analysis

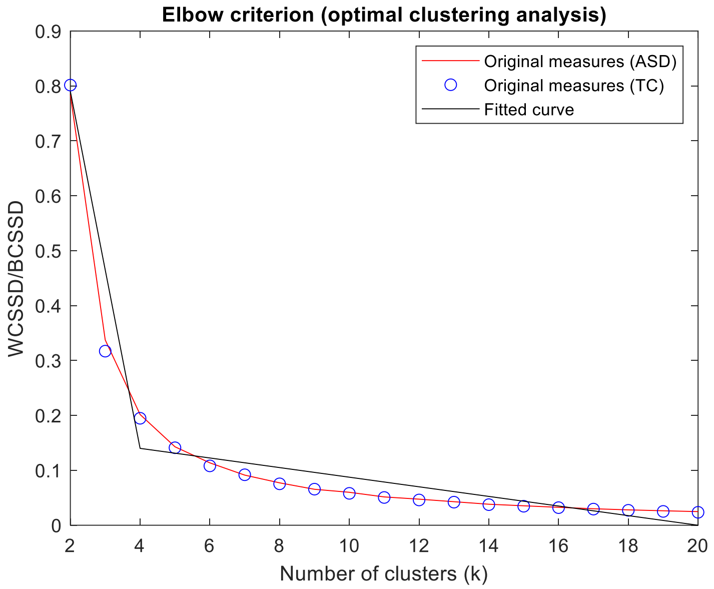


**Figure S1.** The Elbow criterion for selecting the optimum number of clusters of DFC analysis. The red line and blue circles show the original measures of ratio WCSD/BCSSD for ASD and TC with respect to the number of clusters (k), and the black line represents the best fitted elbow-shaped curve to the ASD and TC original measures. The optimum cluster number is 4.
